# Loss of Dicer1 in mouse embryonic fibroblasts impairs ER stress-induced apoptosis

**DOI:** 10.1101/028985

**Authors:** Ananya Gupta, Danielle E. Read, Deepu Oommen, Afshin Samali, Sanjeev Gupta

## Abstract

**Background:** The endoplasmic reticulum (ER) is the site of folding for membrane and secreted proteins. Accumulation of unfolded or misfolded proteins in the ER triggers the unfolded protein response (UPR). The UPR can promote survival by reducing the load of unfolded proteins through upregulation of chaperones and global attenuation of protein synthesis. However, when ER stress is acute or prolonged cells undergo apoptosis. In this study we sought to determine the effect of globally compromised microRNA biogenesis on the UPR and ER stress-induced apoptosis

**Results:** Here we report the role of Dicer-dependent miRNA biogenesis during the UPR and ER stress-induced apoptosis. We show that ER stress-induced caspase activation and apoptosis is attenuated in Dicer deficient fibroblasts. ER stress-mediated induction of GRP78, the key ER resident chaperone, and also HERP, an important component of ER-associated degradation, are significantly increased in Dicer deficient cells. Expression of the BCL-2 family members BIM and MCL1 were significantly higher in Dicer-null fibroblasts. However, ER stress-mediated induction of pro-apoptotic BH3 only protein BIM was compromised in Dicer mutant cells.

**Conclusions:** These observations demonstrate key roles for Dicer in the UPR and implicate miRNAs as critical components of UPR.

## Introduction

The endoplasmic reticulum (ER) is a multifunctional signaling organelle that controls a wide range of cellular processes. The major physiological functions of the ER include folding of membrane and secreted proteins, synthesis of lipids and sterols, and storage of free calcium. Cellular stresses that impair proper folding of proteins can lead to an imbalance between the load of resident and transit proteins in the ER and the ability of the organelle to process that load. In mammals, three ER transmembrane protein sensors, Ire1, Atf6 and Perk, respond to the accumulation of unfolded proteins in the ER lumen [1]. Activation of Perk, Ire1 and Atf6initiates ER-to-nucleus intracellular signaling cascades collectively termed as the unfolded protein response (UPR). The most salient feature of the UPR is to increase the transactivation function of a gamut of bZIP transcription factors, such as Atf6, Atf4 and Xbp1 [2]. Once activated, these transcription factors coordinate transcriptional induction of ER chaperones and genes involved in ER-associated degradation (ERAD) to enhance the protein folding capacity of the cell and to decrease the unfolded protein load of the ER, respectively [1,2]. However, if the damage is too severe and ER homeostasis cannot be restored, apoptosis ensues [3]. The exact mechanism involved in transition of the UPR from protective to pro-apoptotic is not clearly understood.

A class of small RNAs, known as microRNAs (miRNAs), have been shown to be critically involved in control of cell survival and cell death decisions [4-6]. miRNAs are generated from RNA transcripts that are exported into the cytoplasm, where the precursor-miRNA molecules undergo Dicer1-mediated processing to generate mature miRNA [7]. The mature miRNAs assemble into RNA-induced silencing complexes (RISCs) and guide the silencing complex to specific mRNA target molecules with the assistance of argonaute proteins. The main function of miRNAs is to direct posttranscriptional regulation of gene expression, typically by binding to 3’ UTR of cognate mRNAs and inhibiting their translation and/or stability by targeting them for degradation [8].

Several studies have shown global alterations in miRNA-expression profiles during various types of cellular stresses, such as folate deficiency, arsenic exposure, hypoxia, drug treatment and genotoxic stress [9,10]. Argonaute family member Ago2, a vital component of RISCs, is distributed diffusely in the cytoplasm and redistributes from the cytoplasm to stress granules and processing (P)-bodies upon exposure to stress conditions [11]. Stress-induced enrichment of Ago2 from cytoplasm to P-bodies is dependent on mature miRNAs suggesting a link between miRNAs and cellular stress. The miRNAs have emerged as key regulators of ER homeostasis and important players in UPR-dependent signaling [12,13]. We have recently shown that human colon cancer cells hypomorphic for Dicer1 activity are resistant to ER stress-induced apoptosis [14]. In this study we have used global approach to determine the role of miRNAs in ER stress responses by deleting Dicer1, in mouse embryonic fibroblasts. As expected the loss of Dicer1 results in a dramatic decrease in the level of mature miRNAs. We found that Dicer1 lacking fibroblasts were resistant to ER stress-induced apoptosis as compared to wild type (WT) fibroblasts. In addition, loss of Dicer1 enhanced the induction of GRP78 and HERP upon exposure to ER stress. Furthermore, Dicer1-deficient cells show increased basal expression of BIM and MCL1. However, ER stress-mediated induction of pro-apoptotic BH3 family member, BIM is abrogated in Dicer1 deficient cells. These results suggest key roles for miRNAs in cellular responses to ER stress since their global absence compromises ER stress-induced apoptosis.

## Materials and Methods

### Cell culture and treatments

Dicer1 conditional Mouse embryonic fibroblasts (MEFs) were kind gift from Dr. Brian Harfe, University of Florida, Gainesville, USA [15]. MEFs were maintained in Dulbecco’s modified medium (DMEM) supplemented with 10% FCS, 100 U/ml penicillin and 100 mg/ml streptomycin at 37 °C with 5% CO2. To delete Dicer1, Dicer1 conditional MEFs were infected at a high MOI with Ad-CMV-Cre-GFP adenovirus (Vector Biolabs). Cells were treated with thapsigargin and tunicamycin for the times indicated. All reagents were from Sigma-Aldrich unless otherwise stated.

### RNA extraction, RT-PCR and quantitative RT-PCR

Total RNA was isolated using RNeasy kit (Qiagen) according to the manufacturer’s instructions. Reverse transcription (RT) was carried out with 2 μg RNA and OligodT (Invitrogen) using 20 U Superscript II Reverse Transcriptase (Invitrogen). Real-time PCR method to determine the induction of UPR target genes has been described previously [68]. Briefly, cDNA products were mixed with 2 × TaqMan master mixes and 20 × TaqMan Gene Expression Assays (Applied Biosystems) and subjected to 40 cycles of PCR in StepOnePlus instrument (Applied Biosystems). Relative expression was evaluated with ΔΔCT method.

### Analysis of caspase activity

Caspase activity was determined as described previously [16] using carbobenzoxy-Asp-Glu-Val-Asp-7-amino-4-methyl-coumarin (DEVD-AMC). Briefly, assay buffer (100 mM HEPES, 10% sucrose, 0.1% CHAPS, 5 mM DTT, 0.0001% NP-40, pH 7.25) containing 50 μM DEVD-AMC (carbobenzoxy-Asp-Glu-Val-Asp-7-amino-4-methylcoumarin) (Peptide Institute Inc., Japan) was added to samples. Cleavage of the substrate DEVD-AMC to liberate free AMC was measured using a Wallac 1420 Multilabel counter (Perkin Elmer Life Sciences) with 355 nm excitation and 460 nm emission wavelengths over 25 cycles at 1 min intervals.

### Statistical Analysis

The data is expressed as means ± SD for three independent experiments. Differences between the treatment groups were assessed using Two-tailed unpaired student’s t-tests. The values with a *p*<0.005 were considered statistically significant.

## Results

The Dicer1 conditional allele used in this study has the exon 24, encoding one of the RNase III domains flanked by loxP sites (Fig. 1A). Cre-mediated recombination of this exon has been shown to be effective in reducing the ability of Dicer1 to process precursor-miRNAs into mature miRNAs [15]. We infected Dicer1 conditional fibroblasts with an adenovirus expressing the Cre recombinase protein (Ad-Cre-GFP) and assessed inactivation of Dicer1 in MEFs by RT-PCR using primers flanking the floxed exon 24 (Fig. 1A). The majority of Dicer1 transcripts in WT MEFs contained exon 24, however a minimal amount of exon 24 lacking transcripts were also present (lane 1). The presence of exon 24 lacking mRNA may be due to low level of alternative splicing. As expected, two representative Dicer1-deficient clones showed exon 24 lacking mRNA as the major transcript (lane 2, 3). Quantification of mature miRNA levels revealed a significant depletion of mature miRNAs in Dicer1-deficient MEFs, but the extent of depletion differed for individual miRNAs (Fig 1B). Collectively, these results indicate that expression of Cre recombinase in Dicer1 conditional MEFs efficiently depletes Dicer1 activity and compromises biogenesis of mature miRNAs. Loss of Dicer1 has been shown to compromise the proliferative capacity of primary MEFs and embryonic stem cells [17-19]. We observed that Dicer1-deficient cells represented ~90% of the population at passage 2 after infection with Ad-Cre-GFP. However, Dicer1-deficient population was rapidly depleted and at 12 passages post-Ad-Cre-GFP infection the majority of the cells in the population had wild type Dicer1 (Fig 1C and D). Therefore cells between passages 2-6 after infection with Ad-Cre-GFP were used for subsequent experiments.

**Figure 1.**
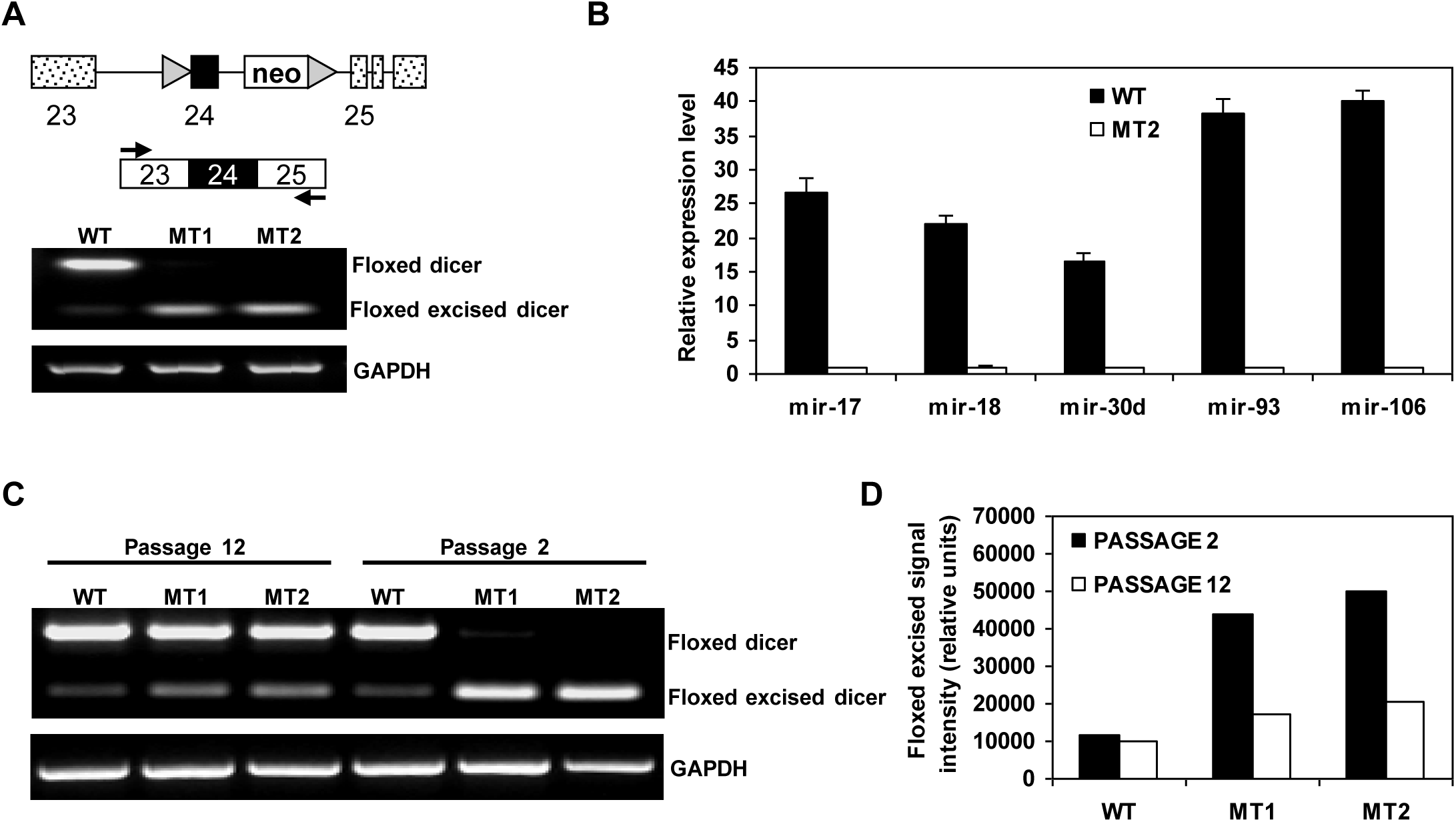
Reduced miRNA biogenesis and proliferation in Dicer1-deleted immortalized MEFs. (A) Schematic representation of Dicer1 conditional allele. *Upper panel*, Dicer1 conditional allele contains loxP sites (triangles) flanking the exon 24 (black box). A neomycin resistance cassette (neo) is retained in intron 24. The positions of primers used for amplifying Dicer1 are indicated by arrows. *Lower panel*, RT-PCR analysis of total RNA from wild type (WT) and Dicer1-deleted (MT1 and MT2) MEFs using primers for Dicer1 and GADPH. (B) Quantification of miRNA levels using TaqMan RT-PCR. Levels of mature miRNAs are normalized to GAPDH expression. Ratios of miRNA to control RNA are plotted in arbitrary units ± SD with average miRNA levels from MT2 set at 1. Results are representative of two different experiments performed in triplicates. (C) Dicer1 floxed MEFs were infected with Ad-Cre-GFP virus and the population was analyzed at different times after Cre expression by RT-PCR. The Dicer1 deficient population was lost from the mixed population at passage 12 after deletion of Dicer1. (D) The relative representation of the Dicer1 deleted cells in the population was quantified by image analysis of the gel shown in C.

Next we investigated the consequences of impaired miRNA biogenesis in Dicer1-null MEFs by determining the extent of cell death after ER stress. ER stress was induced by treatment of cells with either thapsigargin, an inhibitor of the sacroplasmic/Endoplasmic Reticulum Ca^2+^ATPase (Serca) pump, or tunicamycin, an inhibitor of *N*-linked glycosylation. We expressed either Cre-GFP or GFP in Dicer1 conditional MEFs and found that ER stress-induced cell death was attenuated in the Dicer1-deficient, Cre-GFP expressing conditional MEFs (MT2) as compared to GFP expressing conditional MEFs (WT) (Fig 2A). No difference was seen in ER stress-induced cell death in WT MEFs expressing either Cre-GFP or GFP (data not shown). Furthermore, MT2 cells showed reduced DEVDase activity as compared to WT controls (Fig 2B).

**Figure 2.**
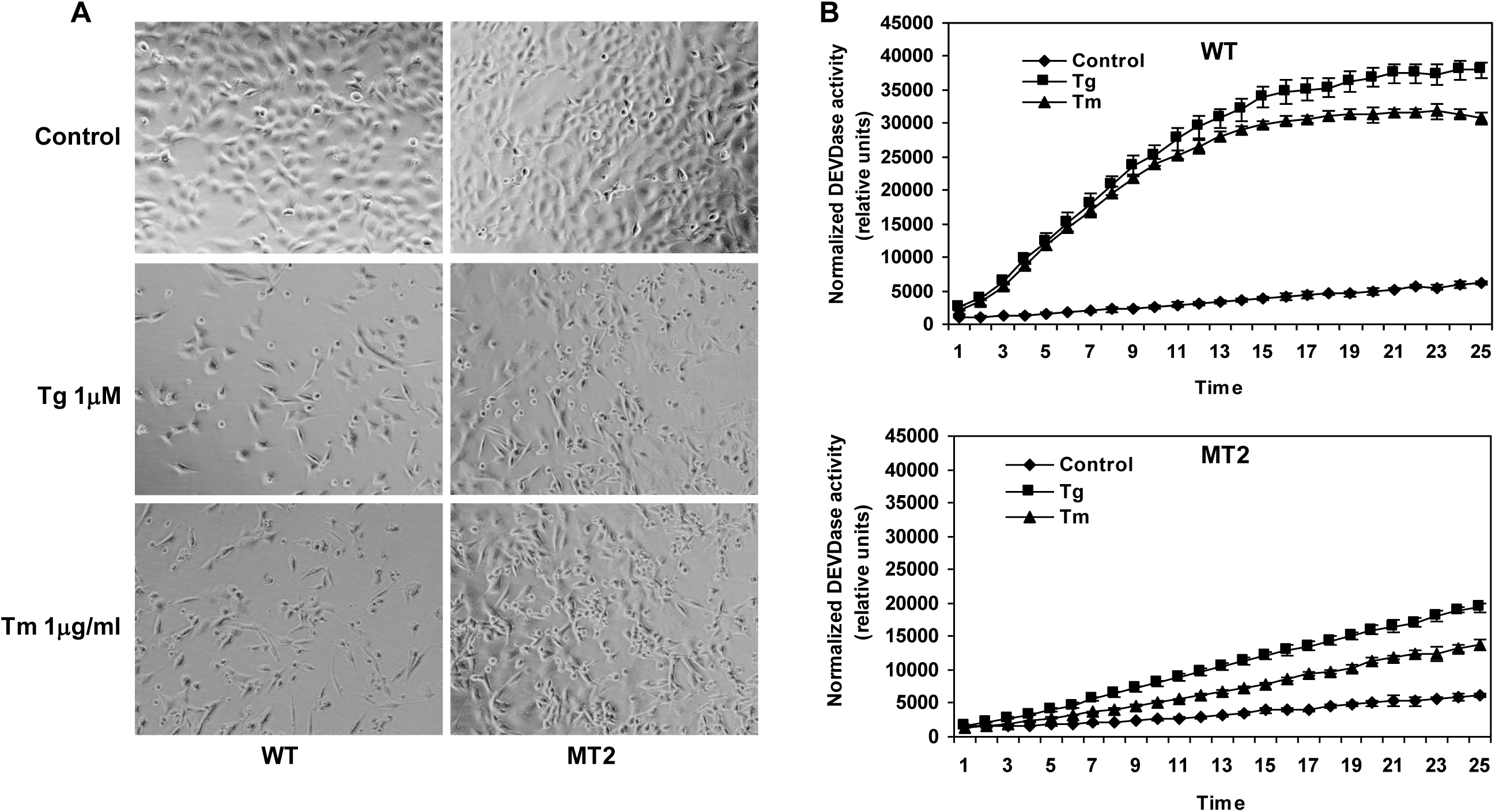
Dicer1-deficient MEFs are resistant to ER stress-induced apoptosis. (A) Wild type (WT) and Dicer1-deleted (MT2) MEFs were either untreated or treated with thapsigargin (1.0 µM) or tunicamycin (1.0 µg/ml) for 24 h. Representative images are shown at 400X magnification. (B) Cells were treated as in A, and DEVD-AMC cleavage activity was measured in whole cell extracts. The DEVDase activity was normalized amount of protein as determined by Bradford assay. Ratios of AMC fluorescence to amount of protein are plotted in arbitrary units ± SD. Values are representative of three separate determinations performed in triplicates.

To investigate the basis for reduced ER stress-induced cell death associated with loss of miRNA biogenesis, we compared the induction of key UPR target genes between WT and Dicer1-deficient MEFs. Quantitative RT-PCR showed that induction of Grp78 and Herp were significantly enhanced in Dicer1-null cells (MT2) as compared to WT cells (Fig 3A). However, there was no difference in the induction of several other UPR target genes such as Chop, Hox1, Edem1, Erp72, p58ipk, Wars and Atf4 (Fig 3A). In the absence of Dicer1, we detected a significant decrease in spliced Xbp1 transcripts upon exposure to ER stress (Fig 3B). Currently, we are unsure of the molecular mechanisms that would account for the downregulation of spliced Xbp1 in the absence of Dicer1. However, indirect effects such as activation of an inhibitor could be one explanation. Quantitative RT-PCR analysis also showed increase in basal expression of Bim and Mcl1 mRNA levels in Dicer1-deficient cells (Fig 3C). The BH3-only proteins Bim, Puma and Noxa are transcriptionally induced by ER stress [20,21] and we found that ER stress-mediated upregulation of Bim was abrogated in Dicer1-deficient cells (Fig 3D). However, Puma and Noxa were induced to a similar extent in both WT and Dicer1 mutant MEFs (Fig 3D). Further we did not observe any significant change in the level of Mcl1 upon exposure to ER stress (Fig 3D). Taken together these results suggest a role for miRNAs in regulating the expression of Bcl2-family members during the conditions of ER stress.

**Figure 3.**
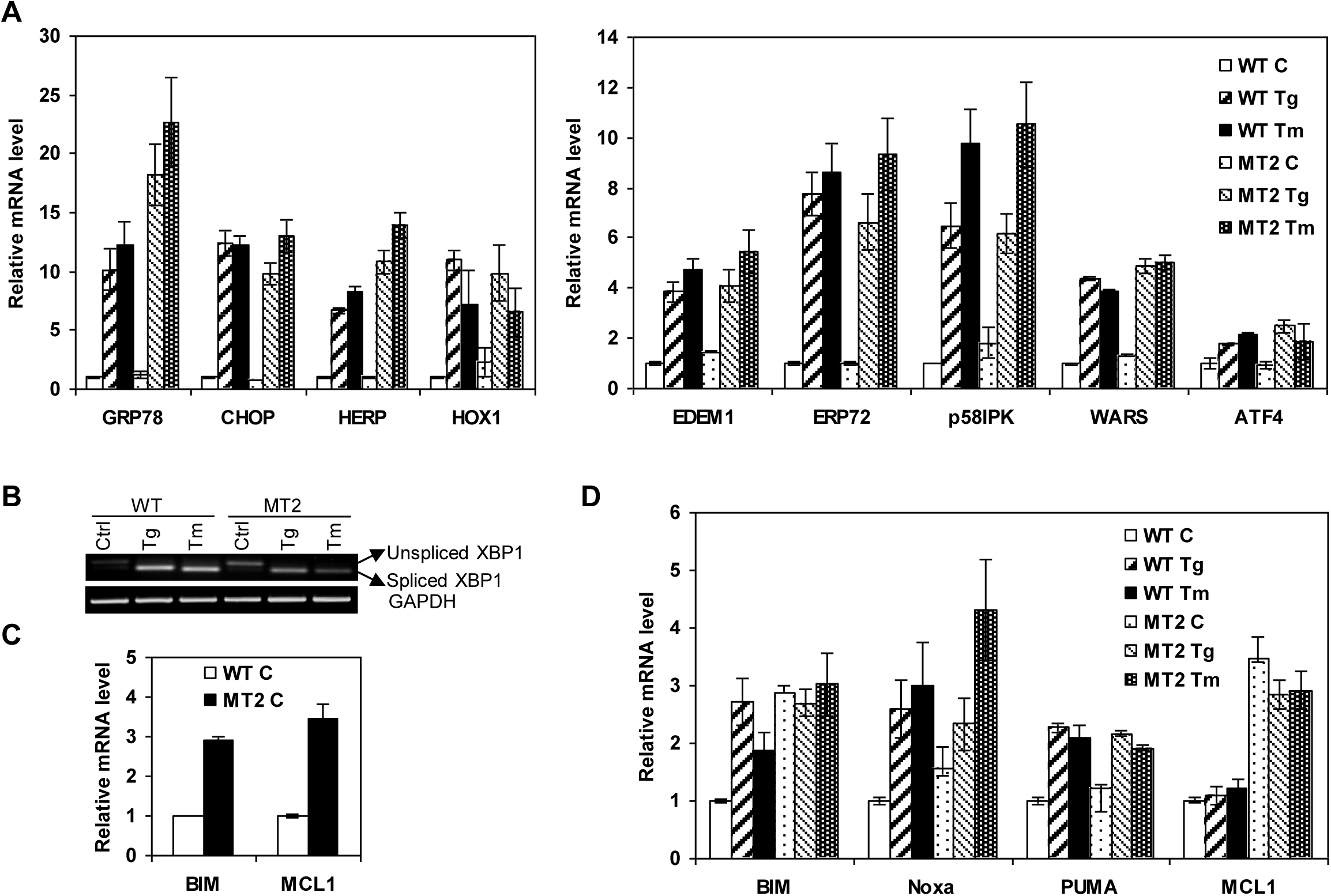
Loss of Dicer1 modulates the induction of specific UPR target genes. (A) Total RNA was isolated from Wild type (WT) and Dicer1-deleted (MT2) MEFs, either untreated or treated with thapsigargin (1.0 µM) or tunicamycin (1.0 µg/ml) for 24 h, and the expression levels of the indicated genes were quantitated by real-time RT-PCR, normalizing against Gapdh expression. Ratios of indicated gene to Gapdh are plotted in arbitrary units ± SD with expression level from wild type, untreated control set at 1. Error bars represent mean ± SD from two experiments performed in duplicate. (B) Wild type (WT) and Dicer1-deleted (MT2) MEFs were treated as in A, and RT-PCR analysis of total RNA was performed to simultaneously detect both spliced and unspliced Xbp1 mRNA and Gapdh. (C) Total RNA was isolated from wild type (WT) and Dicer1-deleted (MT2) MEFs and the expression levels of the indicated genes were quantitated by real-time RT-PCR, normalizing against Gapdh expression. Ratios of indicated gene to Gapdh are plotted in arbitrary units ± SD with expression level from wild type, set at 1. Error bars represent mean ± SD from three experiments performed in duplicate. (D) Wild type (WT) and Dicer1-deleted (MT2) MEFs were treated as in A, and the expression levels of the indicated genes were quantitated by real-time RT-PCR, normalizing against Gapdh expression. Ratios of indicated gene to Gapdh are plotted in arbitrary units ± SD with expression level from wild type, untreated control set at 1. Error bars represent mean ± SD from two experiments performed in duplicate.

## Discussion

In this study we have used conditional Dicer1 MEFs to evaluate the global role of miRNAs during ER stress-induced apoptosis. Our results show that loss of Dicer1 in MEFs leads to a substantial reduction in mature miRNAs and compromises the long term proliferative capacity of the cells. Our work demonstrates that reduced mature miRNA expression can attenuate ER-stress-induced apoptosis. Nevertheless, it is possible that not all of the observed phenotypes are due to the loss of mature miRNAs. It is possible that Dicer1 is processing other dsRNAs or may have nuclear roles as suggested by loss of centromeric and pericentromeric silencing in Dicer1 null ES cells [17,18]. To support this idea, the reduction of DNA methylation has also been associated with Dicer1 loss [22].

Our results show that miRNAs are important players in the ER stress-mediated induction of Grp78, Herp and Bim. The product of each of these genes individually has a known function in regulating ER stress-induced apoptosis. ER chaperone protein Grp78 and ERAD component Herp have been shown to protect cells from apoptosis induced by several insults including ER stress [23-27]. We observed increased basal expression of Bim and Mcl1 in Dicer1 deficient MEFs. Elevated expression of Bim has been reported in Dicer1-deficient pro-B cells and miR-17~92 cluster deficient B cells [28,29]. Furthermore, our group has recently shown that expression of the miR-106b-25 cluster is decreased in a Perk-dependent manner during ER stress which in turn leads to upregulation of Bim and induction of apoptosis [30]. We have also found at least three binding sites for miR-17~92 cluster members in the 3’ UTR of Mcl1 using Targetscan software [31]. Our results showing that expression of members of the miR-17~92 family, such as miR-17, -93 and -106, is significantly downregulated in Dicer1-deficient MEFs as compared to WT MEFs (Fig1B) suggests that decreased expression of the miR-17~92 family contributes to increased expression of Bim and Mcl1 in Dicer1-null fibroblasts and supports emerging data in the literature. Up-regulation of Mcl1 by ER stress is critical for survival of melanoma cells and plays an essential role in antagonizing proapoptotic BH3-only proteins Puma and Noxa [32]. Bim is essential for ER stress-induced apoptosis in a broad range of cell types, including thymocytes, macrophages and epithelial cells [20].

We envision that altered expression of Grp78, Herp, Mcl1 and Bim act in concert, rather than individually, to promote cell survival in Dicer1-null MEFs upon exposure to ER stress. The identification of specific regulatory miRNAs for these mRNAs involved in UPR will provide more insight into the molecular mechanisms underlying this process. Indeed, the role of miRNAs in influencing cell fate during conditions of ER stress is only beginning emerge. Identification of miRNAs involved in the cellular response to ER stress will reveal a valuable level of molecular control and may ultimately enable more avenues for therapeutic intervention. The present study highlights the importance of Dicer1-dependent miRNA biogenesis during the UPR and ER stress-induced apoptosis. These observations demonstrate key roles for Dicer1 in the UPR and implicate miRNAs as critical components of UPR, thus supporting recent findings in the field. The role of miRNA in regulation of the UPR is still an emerging area and further research is required to gain understanding of the regulation involved and to provide additional therapeutic opportunities.

## List of abbreviations

Herp, Hyperhomocysteinemia-induced ER stress-responsive protein; Ire1, Inositol requiring 1; Atf6, Activating transcription factor 6; Mcl1, Myeloid cell leukaemia sequence 1; Perk, Pancreatic eIF2a kinase; Xbp1, X-box-binding protein 1; bZIP, basic region leucine zipper; Grp78, Glucose-regulated protein of 78 kDa; Chop, CCAAT/enhancer-binding protein (C/EBP) homologous protein; Hox1; Haemoxygenase 1; Edem1, ER degradation enhancer, mannosidase a-like 1; Erp72, ER protein of 72 kDa; p58ipk, 58-kDa inhibitor of PKR; Wars, tryptophanyl-tRNA synthetase; Atf4, Activating transcription factor 4; Bim, BCL-2 interacting mediator of cell death; Puma, p53 upregulated modulator of apoptosis; Noxa, Phorbol-12-myristate-13-acetate-induced protein 1; MEF, Mouse embryonic fibroblasts; miRNAs, microRNAs; pri-miRNAs, primary microRNAs

## Acknowledgements

We would like to thank Maria Ryan for invaluable technical assistance. This publication has emanated from research conducted with the financial support of Health Research Board (grant number HRA_HSR/2010/24) to S.G. and Research Frontier’s Programme (RFP) grant number 05/IN3/B851 and Investigator Award number 05/IN3/B851from Science Foundation Ireland (SFI) to A.S.

